# The effect of lifelong physical (in)activity on knee extensor force control

**DOI:** 10.1101/2023.02.28.530405

**Authors:** Jamie Pethick

**Author notes:** **Corresponding author:** Dr Jamie Pethick, School of Sport, Rehabilitation and Exercise Sciences, University of Essex, Wivenhoe Park, Colchester, CO4 3WA, United Kingdom.

## Abstract

It is well-documented that older adults exhibit a greater magnitude and decreased complexity of muscle force fluctuations in comparison to young adults. To date, however, research on this age-related loss of force control has focused on heterogeneous groups of inactive/moderately active older adults, despite accumulating evidence that high levels of lifelong physical activity (such as that exhibited by Masters athletes) has a protective effect on neuromuscular function and morphology. The present study compared healthy young adults (aged < 35; n = 14), healthy but inactive older adults (aged > 55; n = 13) and Masters athletes (aged > 55; n = 14) in order to discern the effects of lifelong physical (in)activity on muscle force control. Force control was assessed during isometric knee extension contractions at 10, 20 and 40% maximal voluntary contraction (MVC) and was quantified according to the magnitude (coefficient of variation [CV]) and complexity (approximate entropy [ApEn]; detrended fluctuation analysis [DFA] α) of force fluctuations. Inactive older adults exhibited significantly greater CV, indicative of poorer force steadiness, than young adults and Masters athletes during contractions at 10, 20 and 40% MVC (all *P* < 0.001). There were no significant differences in CV between the young adults and Masters athletes. These results indicate that lifelong physical activity has a protective effect against the age-related loss of muscle force control and suggest that, up to this point, our understanding of the age-related loss of muscle force control has been confounded by the effects of physical inactivity.

**Key points:**

- Ageing is associated with a decrease in muscle force control (i.e., poorer steadiness and adaptability), though to date this has largely been studied in inactive older adults
- Lifelong physical activity, such as that exhibited by Masters athletes, has a protective role against many age-related decrements in neuromuscular physiology and function
- This study compared force control, during contractions at intensities typical of the requirements of activities of daily living, in healthy young adults, healthy but inactive older adults and age-matched Masters athletes
- Masters athletes exhibited significantly better force steadiness than their inactive counterparts and no difference in steadiness compared to young adults
- Lifelong physical activity appears to modulate the age-related loss of force control, indicating that our current understanding of this loss of force control may be contaminated by the negative effects of inactivity

## Introduction

The ageing neuromuscular system is characterised by a loss of muscle mass and decline in contractile function (Power *et al*., 2013), arising from atrophy and loss of muscle fibres (Lexell *et al*., 1988), changes in motor unit size, properties, and morphology (Piasecki *et al*., 2019), and altered inputs from the nervous system (Castronovo *et al*., 2018). A consequent decreased ability to generate maximal force is important functionally, for activities of daily living (ADLs; e.g., rising from a chair, ascending stairs), and significantly affects quality of life and independence (Wilkinson *et al*., 2018). However, many ADLs (e.g., descending stairs, balance, object manipulation, driving) require only a fraction of maximal force (Kern *et al*., 2001) and instead require the precise control of submaximal force (Lodha *et al*., 2016; Hamilton *et al*., 2019; Davis *et al*., 2020). Indeed, the ability to control force has been suggested to be more closely related to declines in functional capacity than a reduced ability to generate maximal force (Hepple and Rice, 2016) and is a greater predictor of (dys)function (Mani *et al*., 2018). Thus, age-related changes in muscle force control have the potential to have equally, if not greater, detrimental effects on quality of life and independence as the decreased ability to generate force.

Muscle force output is modified according to task demands through the processes of recruitment/derecruitment of motor units, and modulation of their discharge rates (Hepple and Rice, 2016). The resultant force should, ideally, be smooth and accurate, though in reality fluctuates around a prescribed target (Enoka *et al*., 2003). Classically, muscle force fluctuations have been quantified according to their magnitude, using metrics such as the standard deviation (SD) and coefficient of variation (CV; Enoka *et al*., 2003). Such magnitude-based measures provide an index of the degree of deviation from a fixed point within a time-series (i.e., force steadiness) and assume that fluctuations are random and independent (Slifkin and Newell, 1999). However, fluctuations in muscle force are neither random nor independent but rather possess a statistically irregular temporal structure or “complexity” (Slifkin and Newell, 1999). Complexity is typically quantified using metrics such as approximate entropy (ApEn; Pincus, 1991), which quantifies the degree of regularity/randomness in an output, and detrended fluctuation analysis (DFA; Peng *et al*., 1994), which identifies long-range (fractal) correlations and noise colour in an output. These complexity-based measures characterise the moment-to-moment relationship between successive points in a time-series (Pincus, 1991) and reflect the adaptability of force production (Goldberger *et al*., 2002a), i.e., the ability to adapt force output rapidly and accurately in response to task demands (Pethick *et al*., 2016). It has been suggested that the use of both magnitude- and complexity-based measures is necessary to provide a complete understanding of force control (Pethick *et al*., 2021a).

Older adults (aged >60 years) have been repeatedly demonstrated to exhibit a greater magnitude of force fluctuations (i.e., decreased steadiness) compared to young adults (aged 20-30 years; Enoka *et al*., 2003; Oomen and van Dieen, 2017; Pethick *et al*., 2022). This is evident in both the small muscles of the upper limbs, associated with fine force control (Galganski *et al*., 1993) and the large muscles of the lower limbs, associated with balance and locomotion (Tracy, 2007). Older adults have also been demonstrated to exhibit lower complexity of force fluctuations (i.e., decreased adaptability) than their younger counterparts in both the upper (Vaillancourt and Newell, 2003) and lower limbs (Fiogbe *et al*., 2021). This impaired ability to control force is likely to result in a neuromuscular response that is insufficient to withstand a perturbation or adequately compensate when performing a task (Hepple and Rice, 2016). Indeed, decreased force steadiness is predictive of poorer performance in tasks of balance (Davis *et al*., 2020; Mear *et al*., 2023), chair rise time and stair climbing power (Seynnes *et al*, 2005), and reactive driving (Lodha *et al*., 2016).

The effects of ageing on the neuromuscular system and, therefore, the structures and processes underlying muscle force control, are far from uniform amongst older adults (Degens and Korhonen, 2012). There is extensive literature demonstrating that lifestyle factors, in particular physical activity (or a lack thereof), is an important factor in slowing (or accelerating) age-related declines in physiological function (Booth *et al*, 2011; Piasecki *et al*., 2019). However, it has been highlighted that previous research on muscle force control and ageing has failed to account for, or recognise the effect of, older adult’s physical activity status (Pethick and Piasecki, 2022). A recent meta-analysis found that most studies on muscle force control and ageing compared young adults with a heterogenous group of sedentary/moderately active older adults, while several studies excluded physically active older adults (Oomen and van Dieen, 2017). This is of importance as firstly, the age-related loss of muscle force control has important functional implications (Hepple and Rice, 2016) and secondly, there are plausible mechanisms by which lifelong physical activity may attenuate the age-related loss of muscle force control (Pethick and Piasecki, 2022).

Masters athletes, a population comprising adults who have maintained very high levels of physical activity (∼3-6 hours training per week for > 20 years; Piasecki *et al*., 2016), represent the ideal model to study the effects of inherent ageing, as they are unaffected by the deleterious effects of inactivity (Lazarus and Harridge, 2017). Indeed, Masters athletes demonstrate minimised age-related losses of maximal strength (McKendry *et al*., 2018) and balance (Leightley *et al*., 2018) in comparison to age-matched inactive controls. Consequently, it has been recommended that future studies on muscle force control and ageing must account for physical activity status and make comparisons between active and inactive older adults (Pethick and Piasecki, 2022). The aim of the present study was, therefore, to determine the effects of ageing and lifelong physical (in)activity on knee extensor force control. To that end, knee extensor force control, quantified using both magnitude-(CV) and complexity-based measures (ApEn, DFA), was compared in three groups: healthy young adults (aged < 35 years), healthy but inactive older adults and Masters athletes (both aged >55 years).

## Method

### Participants

Statistical power was calculated based on previous research investigating muscle force control and ageing (Vaillancourt and Newell, 2003). These calculations indicated that a total sample size of 39 participants (i.e., n = 13 per group) would be necessary to attain adequate statistical power (*P* < 0.05, power = 0.8; G*Power 3.1, University of Düsseldorf, Germany). To that end, 41 participants provided written informed consent to participate in the study, which was approved by the ethics committee of the University of Essex (Ref. ETH2122-0223), and which adhered to the Declaration of Helsinki. This sample comprised 14 recreationally active young adults (Y; aged <35 years; 10 male, 4 female), 13 older inactive adults (OI; aged >55 years; 10 male, 3 female) and 14 Masters athletes (MA; aged >55 years; 11 male, 3 female), whose characteristics are summarised in Table 1. Both males and females were included as recent evidence has demonstrated no sex differences in muscle force CV among either young adults (Guo *et al*., 2022) or Masters athletes (Piasecki *et al*., 2021). All subjects were classified as “healthy” based on the criteria established by Grieg *et al*. (1994). The suitability of each participant was determined using questionnaires assessing their general health, current physical activity (Recent Physical Activity Questionnaire; Besson *et al*., 2010a) and lifelong physical activity (Historical Adulthood Physical Activity Questionnaire; Besson *et al*., 2010b). Exclusion criteria included any known cardiovascular, respiratory, musculoskeletal, or neurological condition, previous surgery in the lower limb, obesity, use of daily medication and suffering from long-COVID. Inclusion criteria for MA was ∼3-6 hour of training per week for >20 years. On average, the MA had been undertaken 6.2 ± 2.9 hours of training per week for 33.1 ± 9.4 years. For MA who remained competitive (7 out of 14), their age-graded performance (i.e., approximate world-record time for athlete’s age divided by the athlete’s actual time) was 77.9 ± 7.1%, indicating a high level of performance relative to respective age group records and favoured event.

**Table 1.** Participant characteristics and physical activity levels.

| | Young<br>(Y; $n = 14$) | Old inactive<br>(OI; $n = 13$) | Masters<br>athletes (MA;<br>$n = 14$) | *P* value | | |
| --- | --- | --- | --- | --- | --- | --- |
|  |  |  |  | Y vs.<br>OI | Y vs.<br>MA | OI vs.<br>MA |
| Age (years) | 23.9 (4.7) | 61.1 (3.9) | 60.4 (4.4) | **>0.001** | **>0.001** | 1.000 |
| Height (m) | 1.71 (0.09) | 1.69 (0.09) | 1.69 (0.07) | 1.000 | 1.000 | 1.000 |
| Weight (kg) | 72.2 (9.7) | 78.3 (15.6) | 73.0 (11.8) | 0.639 | 1.000 | 0.835 |
| BMI (kg<sup>2</sup>/m) | 20.9 (2.2) | 23.1 (4.3) | 21.5 (3.0) | 0.268 | 1.000 | 0.629 |
| Daily steps* | 12842 (4760) | 7275 (3272) | 13988 (3276) | **0.003** | 1.000 | **>0.001** |
| Sedentary time*<br>(hours/day) | 8.5 (1.3) | 9.5 (2.2) | 7.8 (1.4) | 0.315 | 1.000 | 0.064 |
| Light PA*<br>(hours/day) | 13.8 (0.7) | 14.7 (0.8) | 14.4 (1.1) | 0.050 | 0.235 | 1.000 |
| Moderate-vigorous<br>PA* (hours/day) | 0.6 (0.5) | 0.1 (0.3) | 1.0 (0.4) | 0.027 | 0.060 | **<0.001** |

Data are presented as mean (SD). Abbreviations: BMI, body mass index; PA, physical activity.
\*Indicates measurements with different $n$ to that listed in column titles. In these cases, $n$ was 11 for Y, 11 for OI and 12 for MA. Significant differences are highlighted in bold.

Participants’ body mass and height were measured using calibrated scales and stadiometry, and body mass index (BMI) was calculated using these (Table 1). Current physical activity of participants was determined objectively using Actigraphy; with participants wearing an activPAL4 micro (PAL Technologies, Glasgow, UK) for one week. The activPAL4 unit was placed into a nitrile sleeve and then attached to participants left leg at the midline of the anterior thigh using a Tegaderm film dressing (3M Health Care, St Paul, Minnesota, USA). The nitrile sleeve and Tegaderm dressing provided a waterproof barrier, allowing participants to continue wearing the unit when bathing. From this, the average number of daily steps, sedentary time, light physical activity (<3 METS) and moderate-vigorous physical activity (>3 METS) were calculated over a 5-day period (Table 1). Technical issues (relating to the functioning of the activPAL devices) and medical issues (relating the participants experiencing adverse reactions to either the nitrile sleeves or Tegaderm dressing) meant that this physical activity data was only available for 11 of the Y group, 11 of the O group and 12 of the MA group.

### Maximal strength and force control

Participants were seated in the chair of a Biodex System 4 isokinetic dynamometer (Biodex Medical Systems Inc., Shirley, New York, USA), initialised and calibrated according to the manufacturer’s instructions. Their right leg was attached to the lever arm of the dynamometer, with the seating position adjusted to ensure that the lateral epicondyle of the femur was in line with the axis of rotation of the lever arm. Participants sat with relative hip and knee angles of 85° and 90°, with full extension being 0°. The lower leg was securely attached to the lever arm above the malleoli with a padded Velcro strap, whilst straps secured firmly across both shoulders and the waist prevented any extraneous movement and the use of the hip extensors during the isometric contractions. The isokinetic dynamometer was connected via a custom-built cable to a CED Micro 1401-4 (Cambridge Electronic Design, Cambridge, UK). Data were sampled at 1 kHz and collected in Spike2 (Version 10; Cambridge Electronic Design, Cambridge, UK).

Participants were familiarised with the apparatus and testing procedure by performing a series of practice maximal and targeted submaximal isometric knee extension contractions. After ten minutes rest, the isometric maximal voluntary contraction (MVC) of the knee extensors was assessed. Participants performed a series of 3-second MVCs, each separated by 60-seconds rest. These contractions continued until three consecutive contractions were within 5% of each other. Participants were given a countdown, followed by strong verbal encouragement to maximise their effort. The highest instantaneous force obtained from the three trials within 5% of each other was designated as the MVC force. In the majority of cases, participants achieved values within 5% of each other in the first three contractions performed. In no cases, did it take more than four contractions to achieve three consecutive contractions within 5% of each other.

Ten minutes after the establishment of maximal strength, participants performed a series of targeted isometric knee extensor contractions at 10, 20 and 40% of their MVC, in order to establish muscle force control at intensities typical of those of ADLs (Kern *et al*., 2001). The targets were determined from the highest instantaneous force obtained during the preceding MVCs. Participants performed three contractions at each intensity, with contractions held for 12-seconds and separated by 8-seconds rest. The intensities were performed in a randomised order, with two minutes rest between each intensity. Participants were instructed to match their instantaneous force with a 1mm thick target bar superimposed on a display ∼1m in front of them and were required to continue matching this target for as much of the 12-second contraction as possible.

### Data analysis

Maximal strength was determined as the highest instantaneous force obtained during the MVCs. Measures of the magnitude and complexity of force fluctuations from the force control task were calculated based on the steadiest 5 seconds of each contraction, with MATLAB code identifying the 5 seconds of each contraction with the lowest standard deviation (SD).

The magnitude of force fluctuations in each contraction was measured using the coefficient of variation (CV), which provides a measure of the amount of variability in a time-series normalised to the mean of the time-series, i.e., (SD/mean force) × 100, thus facilitating better comparison between groups differing in strength (e.g., young and old adults; Pethick and Piasecki, 2022).

As recommended by Goldberger *et al*. (2002b), the complexity of force fluctuations was assessed using multiple metrics that quantify different aspects of the time-series. The regularity of force output was determined using approximate entropy (ApEn; Pincus, 1991) and the temporal fractal scaling of force was estimated using detrended fluctuation analysis α (DFA; Peng *et al*., 1994). Sample entropy (SampEn) was also calculated, though as shown in Pethick *et al*. (2015), this does not differ from ApEn when 5,000 data points are used in its calculation. The calculations of ApEn and DFA α (and SampEn) are detailed in Pethick *et al*. (2015). In brief, ApEn was calculated with template length, *m*, set at 2 and the tolerance for accepting matches, *r*, set at 10% of the SD of force output, and DFA α was calculated across time scales (57 boxes ranging from 1250 to 4 data points). ApEn quantifies a continuum from 0 to 2; values close to 0 indicate high regularity and low complexity, while values approaching 2 indicate low regularity and high complexity (Pethick *et al*., 2021). DFA α theoretically ranges from ∼0.5 to ∼1.5. When α = 0.5, each value in a time-series is completely random and independent from previous values (i.e., white noise); when α > 0.5, each value is correlated, to some extent, with previous values. An α of 1.0 is indicative of statistically self-similar fluctuations and long-range correlations (i.e., 1/f or pink noise), while an α of 1.5 is indicative of a smooth output with long-term memory (i.e., Brownian noise; Pethick *et al*., 2021).

### Statistics

Data were analysed using SPSS (version 28; IBM Corporation, USA). Figure were created using JASP (version 0.17.1; University of Amsterdam, Netherlands). All data are presented as means (SD). Data were first tested for normality using the Shapiro-Wilk test. One-way ANOVAs with repeated measures were then used to determine significant differences between participant characteristics (age, height, weight, BMI), physical activity levels (average daily step count, sedentary time, light physical activity, moderate-vigorous physical activity) maximal strength, and force control measures (CV, ApEn, DFA). When main effects were observed, student’s independent samples *t*-tests with Bonferroni-adjusted 95% paired-samples confidence intervals (CIs) were used to identify specific differences. Results were deemed statistically significant when *P* < 0.05.

## Results

### Participant characteristics

Participant characteristics are shown in Table 1. The Y group had *n* = 14 subjects (4 female), the OI group had *n* = 13 (3 female) and the MA group had *n* =14 (3 female). Significant age differences existed between the groups (*F* = 282.834, *P* < 0.001). The OI (95% CIs = 31.2, 41.8 years; *P* = 0.001) and MA (95% CIs = 31.0, 41.3 years; *P* = 0.001) groups were older than Y, though there was no significant difference between OI and MA (95% CIs = −4.9, 5.6 years; *P* = 1.000). No significant differences between the groups were evident for height (*F* = 0.601, *P* = 0.554), weight (*F* = 0.936, *P* = 0.401) or BMI (*F* = 1.615, *P* = 0.212).

### Physical activity levels

Participant physical activity levels are shown in Table 1. Due to technical and medical reasons, *n =* 11 (out of 14) in the Y group, *n* = 11 (out of 13) in the OI group and *n* = and 12 (out of 14) in the MA group. Significant differences existed between the groups for daily steps (*F* = 10.590, *P* < 0.001), moderate-vigorous physical activity (*F =* 14.046, *P* < 0.001) and light physical activity (*F* = 3.385, *P* = 0.047); though for this latter, subsequent post-hoc testing revealed no specific differences. The Y group (95% CIs = 953.9, 10425.5 steps/day; *P* = 0.003) and the MA group (95% CIs = 2077.1, 11349.2 steps/day; *P* < 0.001) both took significantly more daily steps than the OI group (Table 1). There was no significant difference between the Y and MA groups (95% CIs = −5659.5, 3612.7 steps/day; *P* = 1.000). The MA group exhibited greater moderate-vigorous physical activity than the OI group (95% CIs = 0.3, 1.3 hours/day; *P* < 0.001). There were no significant differences between the Y and OI groups (95% CIs =−0.03, 0.9 hours/day; *P* = 0.027) and the Y and MA groups (95% CIs = −0.9, 0.08 hours/day; *P* = 0.060). No significant differences were evident for sedentary time (*F* = 3.065, *P* = 0.61).

### Maximal strength

Significant differences existed between the groups for MVC (*F* = 12.601, *P* < 0.001; Figure 1). The Y group (276.9 [59.5] N·m; 95% CIs = 41.2, 158.7 N·m; *P* < 0.001) and the MA group (235.9 [52.0] N·m; 95% CIs = 0.2, 117.7 N·m; *P* = 0.016) exhibited significantly greater knee extensor MVC than the OI group (176.9 [45.3] N·m). There was no significant difference in MVC between the Y and MA groups (95% CIs = −16.6, 98.6 N·m; *P* = 0.130). The MVC of the MA and OI groups equated to 85.1 and 63.9% of the Y group, respectively.

**Figure 1.**
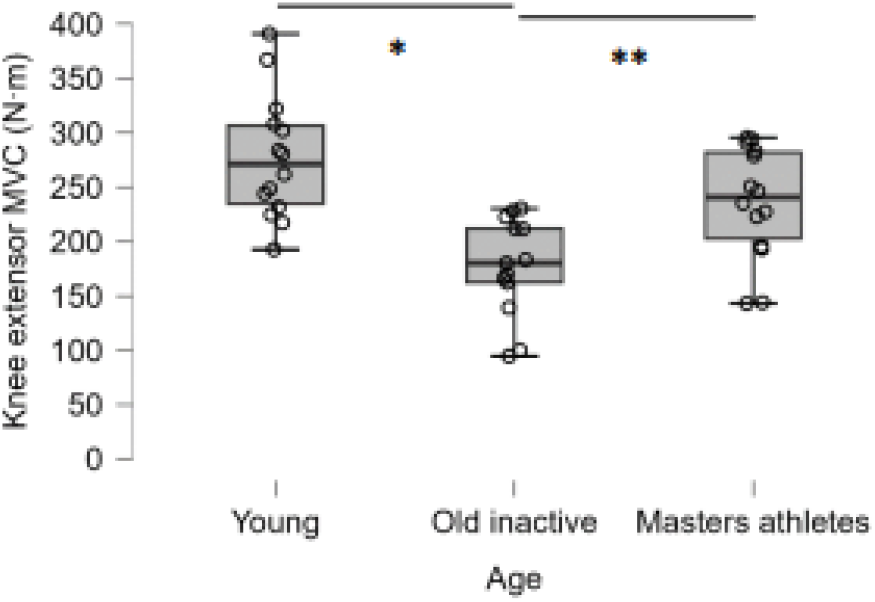
Box and jitter plot of differences in knee extensor maximal strength between groups. * indicates significantly different from Young; ** indicates significantly different from Masters athletes.

### Force control

Significant differences existed between the groups for CV during contractions at 10% (*F* = 17.920, *P* < 0.001), 20% (*F* = 17.741, *P* < 0.001) and 40% MVC (*F* = 12.589, *P* < 0.001; Figure 2; Table 2). The OI group exhibited greater CV than the Y group at 10% (95% CIs = 0.5, 2.1%; *P* < 0.001), 20% (95% CIs = 0.4, 1.5%; *P* < 0.001) and 40% MVC (95% CIs = 0.3, 1.8%; *P* < 0.001). Similarly, the OI group exhibited greater CV than the MA group at 10% (95% CIs = 0.6, 2.2%; *P* < 0.001), 20% (95% CIs = 0.4, 1.4%; *P* < 0.001) and 40% MVC (95% CIs = 0.4, 2.0%; *P* < 0.001; Table 2; Figure 2). There were no significant differences in CV between the Y and MA groups for any contraction intensity (*P* = 1.000 in all cases).

**Figure 2.**
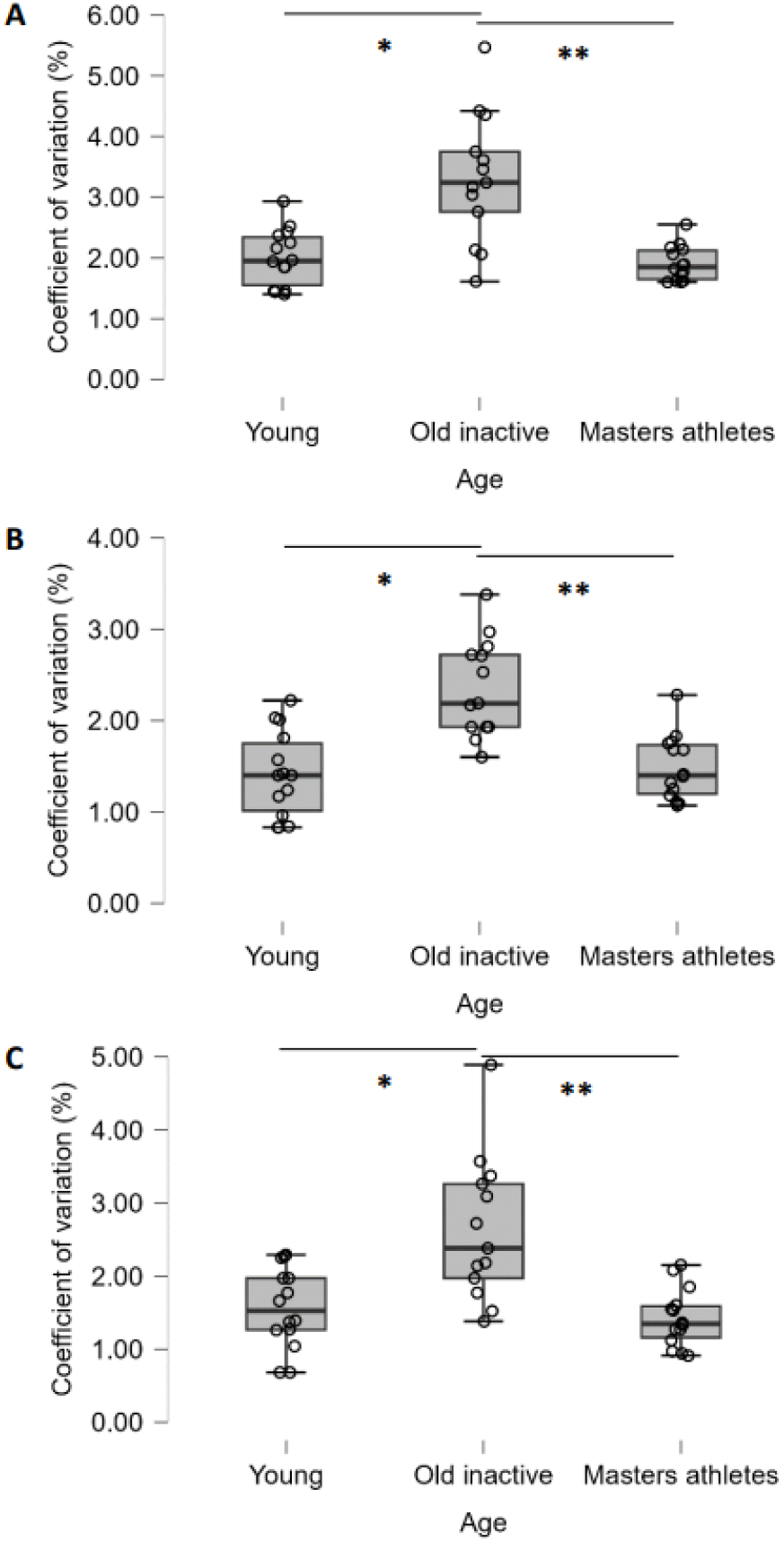
Box and jitter plots of differences in knee extensor coefficient of variation between groups during contractions at 10% (A), 20% (B) and 40% MVC (C). * indicates significantly different from Young; ** indicates significantly different from Masters athletes.

**Table 2.** Force control measures.

|  | Young<br>(Y; *n* = 14) | Old inactive<br>(OI; *n* = 13) | Masters<br>athletes (MA;<br>*n* = 14) | *P* value |  |  |
| --- | --- | --- | --- | --- | --- | --- |
|  |  |  |  | Y vs.<br>OI | Y vs.<br>MA | OI vs.<br>MA |
| CV (%) |  |  |  |  |  |  |
| 10% MVC | 2.00 (0.47) | 3.31 (1.06) | 1.91 (0.29) | **>0.001** | 1.000 | **>0.001** |
| 20% MVC | 1.41 (0.47) | 2.36 (0.53) | 1.49 (0.35) | **>0.001** | 1.000 | **>0.001** |
| 40% MVC | 1.56 (0.55) | 2.63 (0.98) | 1.42 (0.4) | **>0.001** | 1.000 | **>0.001** |
| ApEn |  |  |  |  |  |  |
| 10% MVC | 0.82 (0.05) | 0.81 (0.07) | 0.87 (0.05) | 1.000 | 0.050 | 0.029 |
| 20% MVC | 0.78 (0.07) | 0.77 (0.13) | 0.77 (0.07) | 1.000 | 1.000 | 1.000 |
| 40% MVC | 0.57 (0.09) | 0.56 (0.13) | 0.61 (0.10) | 1.000 | 0.724 | 0.654 |
| DFA $\alpha$ | | | | | | |
| 10% MVC | 1.09 (0.02) | 1.03 (0.10) | 1.06 (0.06) | 0.065 | 0.567 | 0.860 |
| 20% MVC | 1.17 (0.06) | 1.14 (0.11) | 1.16 (0.07) | 0.792 | 1.000 | 1.000 |
| 40% MVC | 1.31 (0.05) | 1.31 (0.08) | 1.25 (0.09) | 1.000 | 0.131 | 0.190 |

Data are presented as mean (SD). Abbreviations: CV, coefficient of variation; ApEn, approximate entropy; DFA, detrended fluctuation analysis. Significant differences are highlighted in bold.

There was a significant effect for ApEn during contractions at 10% MVC (*F* = 4.603, *P* = 0.016), though subsequent post-hoc testing revealed no specific differences (Figure 3; Table 2). No significant differences were evident for ApEn during contractions at 20% (*F* = 0.037, *P* = 0.963) or 40% MVC (*F* = 1.003, *P* = 0.376). No significant differences were evident for DFA α during contractions at 10% (*F* = 2.882, *P* = 0.068), 20% (*F* = 0.643, *P* = 0.531), or 40% MVC (*F* = 2.697, *P* = 0.080; Figure 4; Table 2).

**Figure 3.**
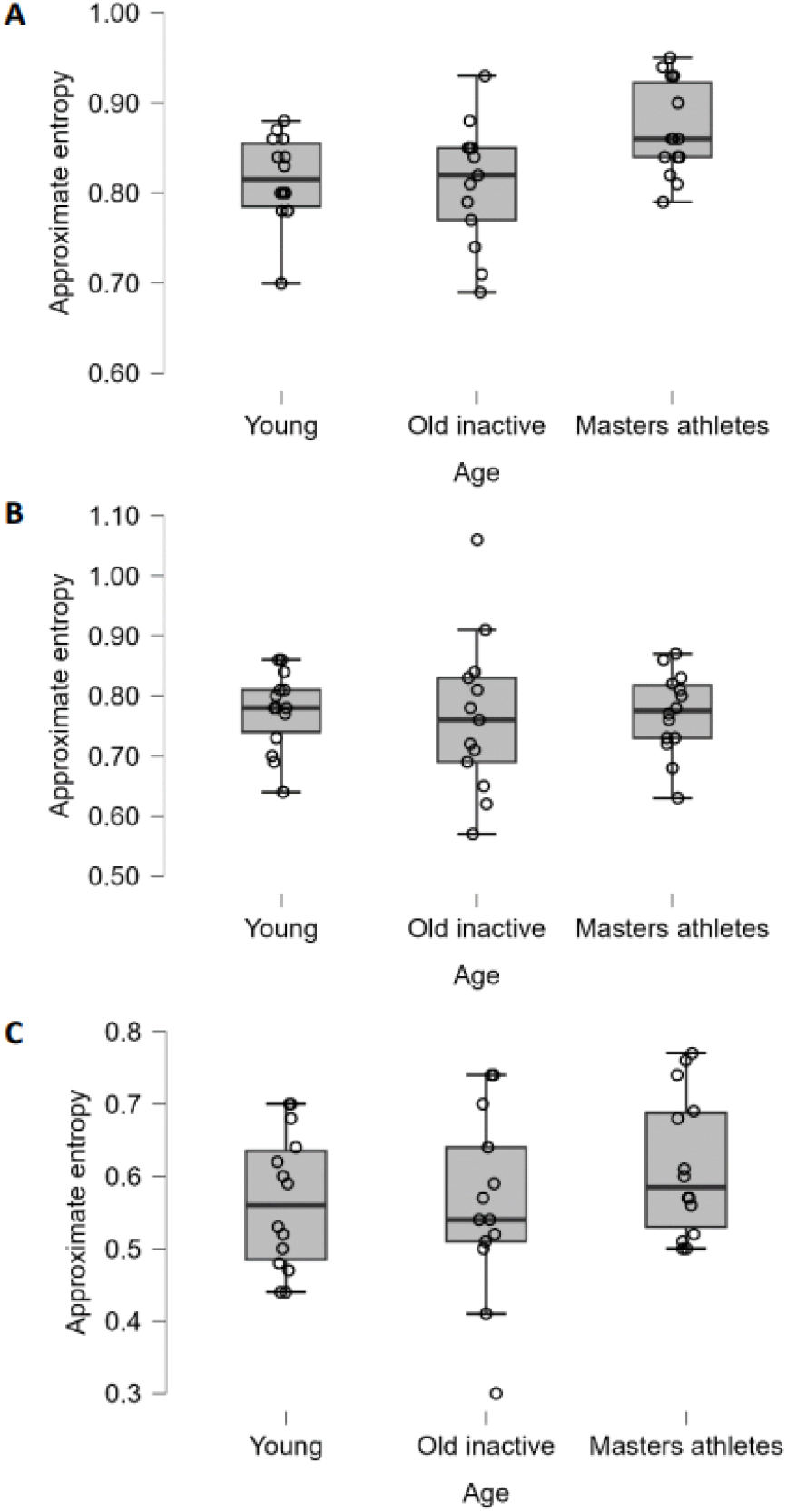
Box and jitter plots of differences in knee extensor approximate entropy between groups during contractions at 10% (A), 20% (B) and 40% MVC (C).

**Figure 4.**
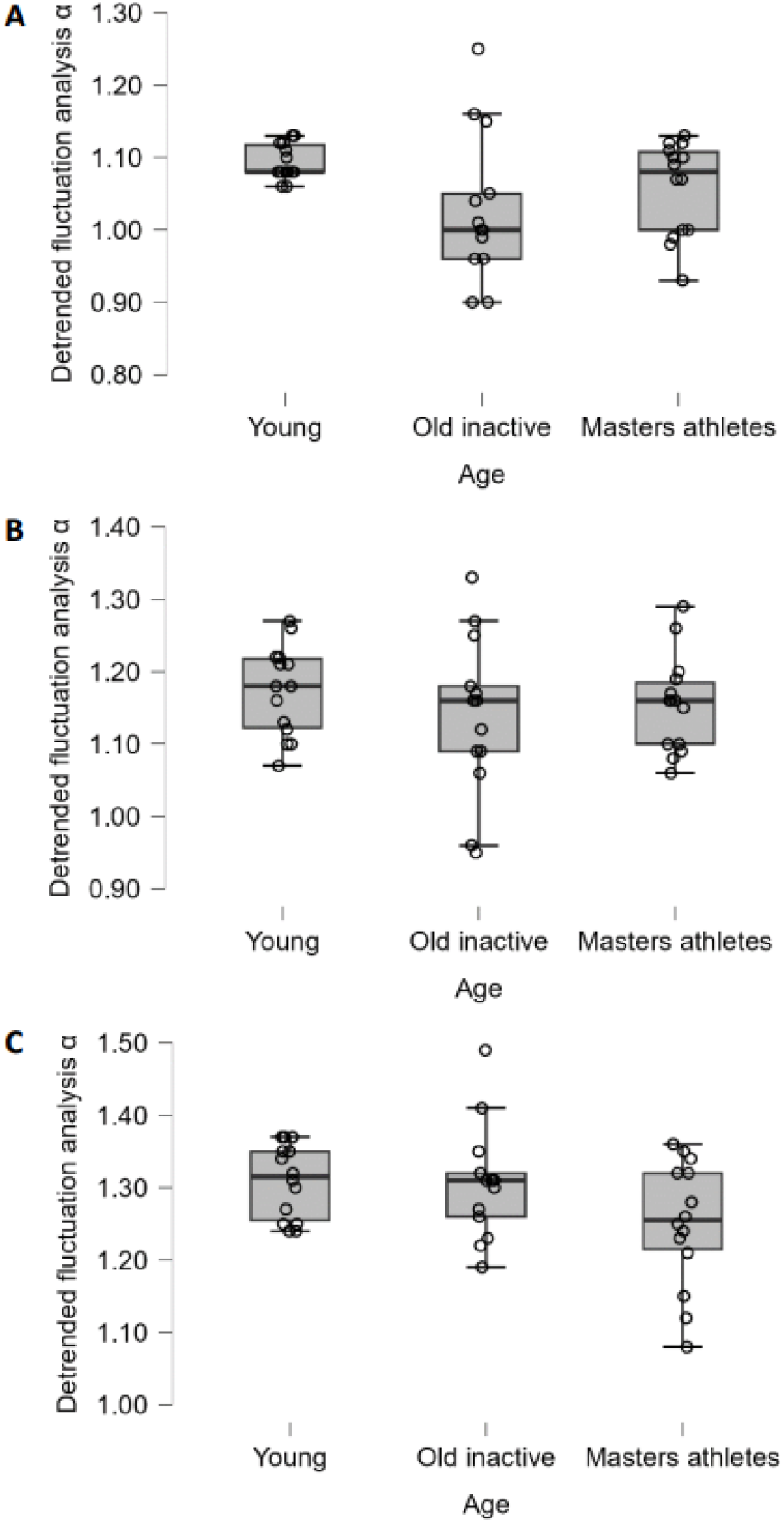
Box and jitters plot of differences in knee extensor detrended fluctuation analysis α between groups during contractions at 10% (A), 20% (B) and 40% MVC (C).

## Discussion

Accumulating evidence has demonstrated differing physiology (e.g., motor unit size) and functionality (e.g., strength) between lifelong physically active and inactive older adults (Piasecki *et al*., 2019). This study is the first to investigate the effect of older adult’s physical activity status on the age-related loss of muscle force control. The major novel finding of the present study was that lifelong physically active older adults (i.e., Masters athletes, aged >55 years) exhibited lower knee extensor force CV, indicative of greater force steadiness, than age-matched inactive adults. Moreover, there were no differences in knee extensor force CV between Masters athletes and healthy young adults. These results provide the first evidence that lifelong physical activity has a protective effect against the age-related loss of muscle force control and further evidence that physical activity status should be considered when investigating the inherent physiological effects of ageing (Lazarus and Harridge, 2010).

### Magnitude-based measures of force control

There is a substantial body of literature demonstrating that older adults exhibit greater muscle force CV (i.e., decreased steadiness) compared with young adults (Enoka *et al*., 2003; Pethick *et al*., 2022), with the knee extensors being one of the most affected muscle groups (Oomen and van Dieen, 2017). The present results are in accordance with these previous observations; inactive older adults exhibited significantly greater knee extensor force CV than young adults during contractions at 10, 20 and 40% MVC (Figure 2; Table 2). Of greater importance, though, was the observation that lifelong physically active older adults (i.e., Masters athletes) exhibited significantly lower knee extensor force CV than the inactive older adults during contractions at 10, 20 and 40% MVC. Indeed, no differences in knee extensor force CV were evident between the Masters athletes and the young adults. This study, therefore, vindicates the assertion that lifelong physical activity has a modulatory effect on the age-related loss of muscle force control (Pethick and Piasecki, 2022). Moreover, these results suggest that the age-related loss of muscle force control previously described (Enoka *et al*., 2003; Oomen and van Dieen, 2017; Pethick *et al*., 2022) is, at least partly, attributable to the deleterious effects of inactivity on the neuromuscular system.

Unfortunately, no electromyographic (EMG) data was obtained in the present study, which could have provided mechanistic insight into the observed differences in muscle force CV. Nevertheless, differences in muscle force CV between populations are indicative of differences in common modulation of motor unit discharge rates and neural drive to muscle (Enoka and Farina, 2021). Recently, Castronovo *et al*. (2018) demonstrated an increase in the common modulation of motor unit discharge rates with age that was highly coherent with the age-related increase in muscle force CV. The present results could, therefore, be indicative of lower common modulation of motor unit discharge rates, and, consequently, lower neural drive to achieve a given relative target force in Masters athletes compared to inactive older adults. Studies on acute physical activity interventions support this supposition. Six-week resistance training programs have been demonstrated to decrease muscle force CV (i.e., increase steadiness) in both young (Vila-Cha and Falla, 2016) and old adults (Kornatz *et al*., 2005), with these changes correlated with, and partially mediated by, decreased motor unit discharge rate variability. To provide further insight into differences in muscle force CV between lifelong physically active and inactive older adults, future research must simultaneously measure muscle force output and motor unit spike trains (using high-density EMG) in these distinct populations. From such high-density EMG measurements, it would also be possible to derive the ApEn of the cumulative motor unit spike train, which has been speculated to be of greater utility than the ApEn of muscle force (Dideriksen *et al*., 2022).

Whilst it seems reasonable to suggest that common modulation of motor unit discharge rates may vary between lifelong physically active and inactive older adults, the sources that contribute to such variance are unknown (Castronovo *et al*., 2018) and elucidating these should be another key goal of future research. Common synaptic input, which drives common modulation of motor unit discharge rates, arises from afferent feedback, descending cortical and reticulospinal pathways, and neuromodulatory pathways from the brain steam (Enoka and Farina, 2021). As discussed in Pethick and Piasecki (2022), there are plausible mechanisms by which lifelong physical activity may influence each of these and, in turn, modulate the age-related loss of muscle force control. In addition to neural drive, force fluctuations are also influenced by the number of motor units in a population (Hamilton *et al*., 2004). Both Masters athletes and inactive older adults exhibit an age-related decrease in motor unit number, though Masters athletes appear to have a greater remodelling capacity via compensatory expansion and re-innervation of denervated muscle fibres (Piasecki *et al*., 2019). Further research is needed to establish what, if any, effect this motor unit expansion has on force control.

### Complexity-based measures of force control

Though under-investigated in comparison to magnitude-based measures of force control, older adults have also been demonstrated to exhibit lower complexity (e.g., ApEn, SampEn, DFA α) than young adults in a range of muscle groups (Vaillancourt and Newell, 2003; Challis, 2006), including the knee extensors (Fiogbe *et al*., 2021). The present results contrast with these previous observations, in that neither ApEn (Figure 3) nor DFA α (Figure 4) differed between the young adults, inactive older adults and Masters athletes, indicating similar levels of force adaptability across the three groups. The reason for this discrepancy with previous results is unknown and further research is warranted to resolve this issue. Nevertheless, the presently observed lack of differences in complexity-based measures between the three groups should not discourage authors from using them. Magnitude- and complexity-based measures provide distinct information and the use of both types of metric will ensure a thorough understanding of age- and (in)activity-related changes in force control (Pethick *et al*., 2021; Clark *et al*., 2023).

### Implications

Muscle force CV during submaximal contractions can explain moderate amounts of variance, often more so than maximal strength, in performance of ADLs involving lower limb muscles, such as static (Davis *et al*., 2020) and dynamic balance (Mear *et al*., 2023), stair climbing power (Seynnes *et al*., 2005) and reactive driving (Lodha *et al*., 2016). Seemingly few studies have investigated performance of ADLs between Masters athletes and inactive older adults, though there is evidence that static (Sundstrup *et al*., 2010; Leightley *et al*., 2017), dynamic (Lee *et al*., 2021) and reactive balance (Brauer *et al*., 2008) are superior in Masters athletes compared to age-matched inactive adults. Given the predictive power of muscle force CV, the presently observed differences between Masters athletes and inactive older adults could contribute to the superior balance others have observed in Masters athletes. Indeed, in young adults a four-week training intervention has been demonstrated to decrease both the magnitude of force fluctuations and postural sway centre of pressure displacements during quiet standing (Oshita and Yano, 2011).

### Conclusion

Masters athletes exhibit greater force steadiness than age-matched inactive adults and similar force steadiness to healthy young adults. This is the first study to demonstrate that lifelong physical activity modulates the age-related loss of muscle force control. These results indicate that our understanding of the age-related loss of muscle force control has, up to this point, been confounded by the pathophysiological consequences of inactivity and emphasise the need for future research on force control to consider the physical activity status of older adults.

